# Genetic diversity of the enteroviruses detected from Cerebrospinal fluid (CSF) samples of patients with suspected aseptic meningitis in northern West Bank, Palestine in 2017

**DOI:** 10.1101/383554

**Authors:** Kamal Dumaidi, Amer Al-Jawabreh, Fekri Samarah, Areej Zreeq, Dirgham Yaseen

## Background

Human enterovirus (HEV) is a non-enveloped RNA genus of the family *Picornaviridae* with more than 107 genotypes. HEV is divided into 4 HEV species designated HEVA to D, based on the sequences analysis of the partial or complete VP1 region of the viral genome[1]. The VP1 are immunodominant capsid proteins that contain the neutralization antigenic determinants which correlate well with serotyping and are used for HEV genotyping[2].

HEVs are associated with several diverse clinical manifestations including mild febrile illness, gastroenteritis, respiratory tract infection, neonatal sepsis-like illness and aseptic meningitis as the most frequent infections caused by HEV that occurs as sporadic and/or outbreaks of varying size. HEV-B causing aseptic meningitis include echoviruses, some enteroviruses coxsachievirus B[3-5]. Furthermore, a sever and potentially fatal conditions including encephalitis, acute flaccid paralysis, myocarditis and hand-foot-mouth disease had been reported [6-10].

In the last two decades, echovirus18 (E18) and coxsackievirus B5 (CVB5), serotypes of HEV-B, have been isolated from sporadic and outbreak cases of aseptic meningitis in China, Taiwan, Germany, Korea, Australia and Palestine. These isolates were found to be associated with various degrees of illness among newborns, infants and school-age children, ranging from asymptomatic to fatal aseptic meningitis [10-17].

The high mutation rate and the RNA recombination are responsible for genetic diversity of the enteroviruses [18]. The recombination increases viral pathogenicity, eliminates lethal mutations and increases fitness of virus [18]. Of the four species, A, B, C and D, HEV-B was shown to have the highest rates of recombination particularly between members of the same species [19-20]. The few studies that investigated the molecular characterization of the two HEVs (E18 and CVB5), showed a wide range of genetic diversity in the VP1 region coding for the outer surface protein, which involved in virus-cell interactions that might explain their high endemicity and infection severity among the newborns, infants, children and adults[12-16]. The aim of the present study was to identify the most predominant enteroviruses which circulated in the northern West Bank, Palestine in 2017 using RT-PCR followed by sequencing and to investigate the genetic diversity of the sequences of the partial VP1 region.

## Materials and Methods

### Sample and data collection

During the period of March, 1^st^ to October, 31^th^, 2017, 249 cerebrospinal fluid (CSF) specimens were collected from children admitted, with suspected aseptic meningitis, to Rafeedia governmental hospital in Nablus and Jenin governmental Hospital in Jenin and were store at -20°C until use. All CSF samples were proven negative for classical bacterial pathogens by the hospital microbiology laboratory. Patients’ demographic data and clinical history including age, sex, sign and symptoms, place of residence, date of onset of symptoms, and CSF laboratory test results were retrieved from the patients’ files. The study was approved by the Ministry of Health in Palestine under reference number ATM/125/2013

### Viral RNA extraction

The HEV RNA was extracted from 200 μl of CSF using a QIAamp viral RNA Mini Kit (QIAGEN, Valencia, CA, USA) according to the manufacturer’s instructions

### RT-PCR

To identify the enterovirus associated aseptic meningitis cases, the HEV RNA was amplified using two primer sets targeting the 5’NCR of the HEV genome as described and modified previously[10, 21]. For HEV genotyping, an RT-PCR with two primer sets was used to amplify the 5’ half of the VP1region of the viral genome as described previously[10, 22]. PCR amplicons were purified using the NucleoSpin Gel and PCR Clean-up from Machery Nagel (Germany) before sending for commercial sequencing.

### Enterovirus typing and phylogenetic analysis

The HEV identity was investigated by searching the partial VP1 sequences obtained in this study in comparison with the HEV prototypes and other HEVs sequences available in GenBank using basic local alignment (BLAST, http://www.ncbi.nlm.nih.gov/BLAST). The HEV that showed more than 75% nucleotide similarity was assigned to be the same genotype. Phylogenetic analyses of the RNA sequences of E18 and CVB5 genotypes were conducted based on the neighbor-joining (NJ) methods with Tamura-Nei and Kimura 2-parameter models implemented in MEGA 6 with 1000 bootstraps replicates for branch confidence in clades in each tree (Tamura et al., 2013). Poliovirus was used as an out-group.

### Recombination analysis

The aligned E18 and CVB5 sequences were tested for any possible recombination events using the RDP 4 software [23]. All seven default statistical tools were employed including PHI statistic (Φ_w_. or pairwise homoplasy index, PHI)[24], Geneconv [25], Bootscan [26], Max X^2^ [27], Chimaera [28], SiScan, and 3Seq [29]. P-value differs depending on the recombination test used, number of sequences analyzed where P-value is subsequently corrected by the RDP 4 default Bonferroni-correction. The recombination tests were performed using default settings and a Bonferroni-corrected *P* value cutoff of 0.05. The recombination analyses were confirmed by SplitTree 4 (ver. 4.14.6) software [30] using PHI statistic [24] and DnaSP 5.1 software [31].

### Genetic diversity analysis and neutrality tests

To account for the limited number of genotype HEV cases (26) and the intermediate length of VP1 sequence (400bp) and the diversity that might be caused external factors such as amplifications and sequencing, several diversity indices were used [32]. The genetic diversity of the 13 E18 and the 11 CVB5 (9 in the present study and the 2 previously detected in 2013) detected in Palestine were analyzed based on VP1 gene. The 27 E18 from 10 different countries reported in the period of 1999-2017 and 25 CVB5 from 13 different countries reported in the period between 1971 and 2017 were retrieved from the Genebank and included in the analysis. EV18 and CVB5 RNA sequences were separately aligned using MEGA 6 [33]. Then, the genetic diversity of the VP1 gene in both E18 and CVB5 genotypes were calculated based on parameters including haplotype diversity (H_d_), nucleotide diversity (π). Haplotype diversity (or gene diversity) refers to the number of haplotypes in the population. Nucleotide diversity is the average number of nucleotide differences per site in pairwise comparison between RNA sequences[34]. To detect departure from neutral theory of evolution (random mutation) at a constant population size, two neutrality tests were used. The first was Tajima’s *D* test which compares the differences between the numbers of segregating sites (S) and the average number of nucleotide differences between two randomly chosen sequences from within in the population (K)[35]. The second test was Fu and Li’s *F* test which compares differences between the number of singleton mutations and the average number of nucleotide differences between pairs of sequences [36]. DnaSP 5.1 software was used for the calculation at default settings[37].

### Statistical analysis

Epidemiologic data were analyzed using EpiInfo statistical package. Analysis included distribution, 2×2 tables, and frequency tables. Fisher’s exact test and Chi square with 95% confidence interval were calculated. The level of statistical significance was P<0.05.

## Results

### Demography of circulating EVs

A total of 249 CSF samples were collected from hospitalized patients suspected of having aseptic meningitis in the period between March-- October, 2017. Overall, 54/249 (22%) yielded positive results for HEV using RT-PCR targeting the 5’UTR. The general characteristic and the clinical history of the study samples are shown in table 1.

**Table 1.** Demographic data and clinical history of the study samples

|  | **Total 249**<br>**No. (%)** | **EV**<br>**54** | **Non-EV**<br>**195** |
| --- | --- | --- | --- |
| Sex: |  |  |  |
| Male | 156 (62.5) | 33 | 123 |
| Female | 93 (27.5) | 21 | 72 |
| Age in month, median (range) | 6 (0-92) | 7 (0-92) | 6 (0-90) |
| Place of residence: |  |  |  |
| Nablus | 156 (62.6) | 26 (48.1) | 130 (66.7) |
| Jenin | 93 (37.4) | 28 (51.9) | 65 (33.3) |
| Pleocytosis | 50/179 (28) | 13 (26) | 37 (74) |
| Fever | 233/249 (93.6) | 179 (77) | 54 (23) |
| Headache | 66/249 (26.5) | 33 (50) | 33(60) |
| Diarrhea | 53/249 (21.3) | 20/54 (37)* | 33/195 |
| Vomiting and Nausea | 82/249 (32.9) | 26/54* | 56/195 |
| Stiff neck | 18/249 (7.2) | 13/41* | 5/195 |
| Photophobia | 9/249 (3.6) | 7/54 (13)* | 2/195 |
| Mental disturbances | 5/249 (2) | 2 (40) | 3(60) |

\* P < 0.05

Twenty-six (48%) were successfully genotyped by sequencing the amplicon of the amplified 5’ VP1 region. Four different types of HEVs were detected. All of them belong to HEV-B species. Thirteen of the detected HEVs (50%) were E18, 9 (34.5%) were coxsackievirus B5 (CVB5), 3 (11.5%) were E25 and 1 (3.8%) was CVB2. Demographic data, the clinical history and the gene Bank accession number of the genotypes HEV cases are shown in Table 2

**Table 2.** Demographic data and clinical characteristic of the diagnosed HEVs genotypes

| Sample No. | Accession No. | EV genotype | Month/year of isolation | Age (month) | District |
| --- | --- | --- | --- | --- | --- |
| 170J/017 | MG773498 | E18 | 04/2017 | 33 | Jenin |
| 186J/017 | MG773499 | E18 | 05/2017 | 64 | Jenin |
| 188J/017 | MG773500 | E18 | 05/2017 | 16.5 | Jenin |
| 208N/017 | MG773501 | E18 | 05/2017 | 70 | Nablus |
| 227J/017 | MG773502 | E18 | 07/2017 | 57 | Jenin |
| 232J/017 | MG773503 | E18 | 08/2017 | 3 | Jenin |
| 247J/017 | MG773504 | E18 | 07/2017 | 5 | Jenin |
| 257J/017 | MG773505 | E18 | 08/2017 | 81 | Jenin |
| 262J/017 | MG773506 | E18 | 08/0217 | 67 | Jenin |
| 265J/017 | MG773507 | E18 | 08/2017 | 3 | Jenin |
| 344J/017 | MG773508 | E18 | 04/2017 | 6 | Jenin |
| 376J/017 | MG773509 | E18 | 09/2017 | 77 | Jenin |
| 380J/017 | MG773510 | E18 | 08/2017 | 10.5 | Jenin |
| 206N/017 | MG773497 | CVB5 | 05/2017 | 40 | Nablus |
| 230J/017 | MG773489 | CVB5 | 07/2017 | 75.5 | Jenin |
| 211N/017 | MG773511 | CVB2 | 05/2017 | <1 | Nablus |
| 260J/017 | MG773490 | CVB5 | 06/2017 | <1 | Jenin |
| 338N/017 | MG773491 | CVB5 | 04/2017 | 4 | Nablus |
| 342J/017 | MG773492 | CVB5 | 05/2017 | 1 | Jenin |
| 345J/017 | MG773495 | CVB5 | 05/2017 | 21 | Jenin |
| 354N/017 | MG773496 | CVB5 | 04/2017 | 35 | Nablus |
| 407N/017 | MG773494 | CVB5 | 03/2017 | 1 | Nablus |
| 433N/017 | MG773493 | CVB5 | 04/2017 | 49 | Nablus |
| 261/017 | MG773514 | E25 | 05/2017 | 42 | Jenin |
| 299N/017 | MG773512 | E25 | 06/2017 | 6 | Nablus |
| 389J/017 | MG773513 | E25 | 09/2017 | 51 | Jenin |

### Phylogenetic analysis of the partial VP1 gene of the E18 and CVB5

Phylogenetic analysis of the partial VP1 gene was conducted using the thirteen E18 and eleven CVB5 strains from Palestine (9 in 2017 and 2 in 2013). Twenty-seven E18 and 25 CVB5 sequences were retrieved from GenBank were included for comparison. The phylogenetic tree for E18 showed three main clusters with all Palestinian isolates uniquely clustering together along with those from China. Similarly, The CVB5 isolates were distributed into three clusters with Palestinian isolates in 2017 clustering together, along with isolates from different areas of the world, whereas, the two Palestinian isolates in 2013grouped within the Predominantly-Chinese cluster (Figure. 1)

**Figure 1.**
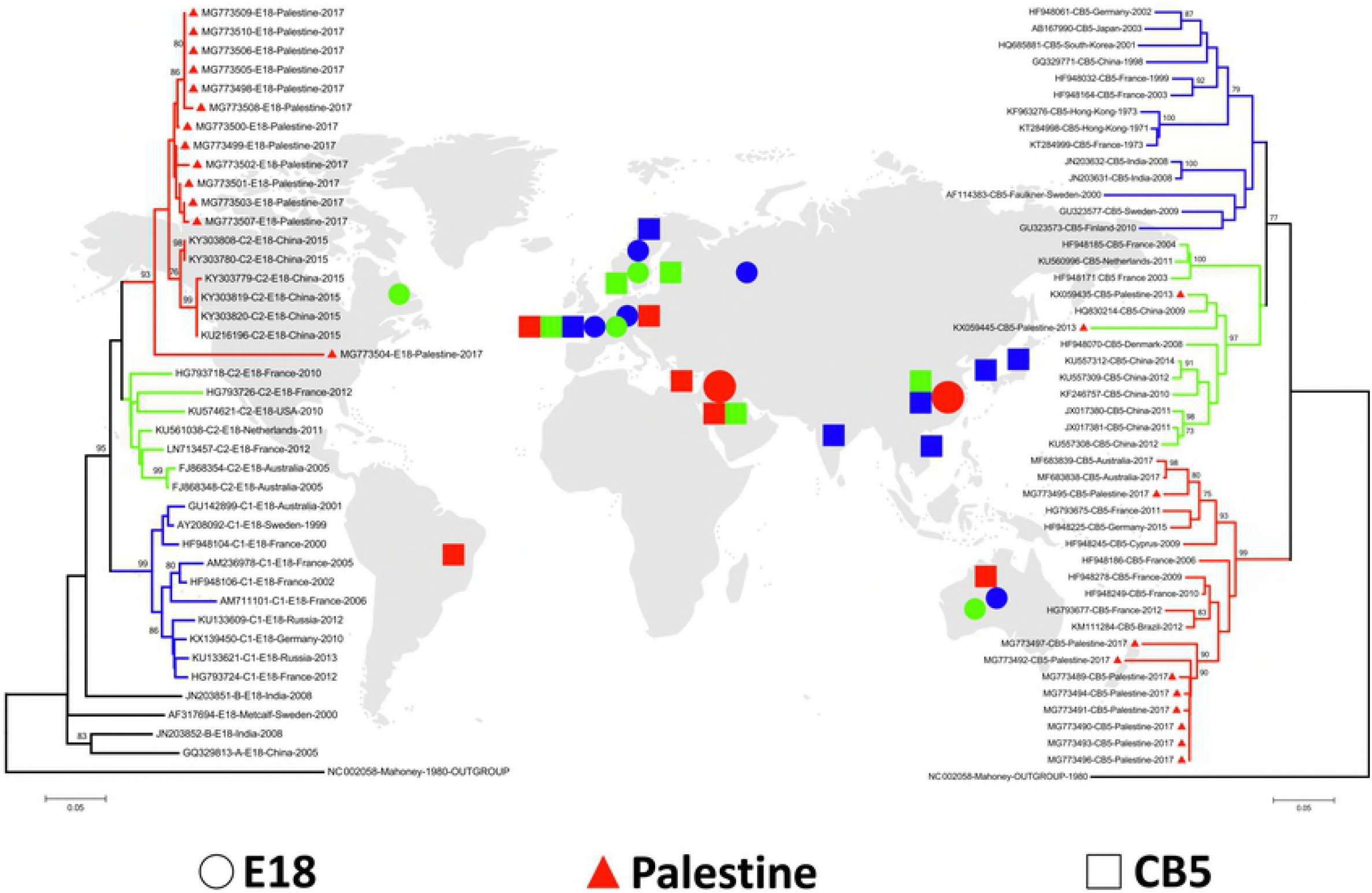
Bootstrap consensus (1000 replicates) neighbor joining (NJ) phylogenetic dendrograms constructed based on the partial VP1 gene of E18 (left) and CVB5 (right). Dendrograms for E18 (o) and CVB5 (□) are plotted against the global distribution of the geographical origin of the isolates. The two cluster contain strains from Palestine (Δ) along with strains from GenBank. Poliovirus from vaccine was included as an out-group.

### Recombination analysis

The recombination analysis did not find any statistically significant evidence for recombination events between the aligned sequences, both on the Palestinian strains level (13 E18 strains and 11 CVB5) and international level that included Palestinian strains pooled with strains from the Gene bank (40E18 strains and 36CVB5). However, minimum number of recombination events was detected by DnaSP 5.1 software for E18 and CVB5. RDP4 software (PHI statistic) revealed good chance of recombination, but at a very low P-value (0.00001).

### Diversity indices

Population nucleotide diversity indices and neutrality tests were calculated for the partial VP1 gene for E18 and CVB5, separately, based on phylogenetic clusters (Tables 3 and 4). The totally haplotype diversity (Hd) for the 40 E18 sequences was 0.98± 0.01 and 0.99± 0.006 for the 36CVB5. At the same time, the total genetic diversity (π) for E18 was 0.12± 0.02 and 0.17± 0.01 for CVB5. The average number of nucleotide differences between any two sequences (k) for E18 and CVB5 were 35.5 and 41, respectively. Cluster I of E18 showed peculiar results compared to the other two clusters. In this group, we detected 12 haplotypes in 19 sequences with the lowest haplotype-to-sequence (h:n) ratio (0.6:1), compared to equal h:n ratio for clusters II and III. Haplotype diversity (H_d_), nucleotide diversity (π), and number of segregating (polymorphic) sites (S) for E18 were lowest in cluster I which is comprised mainly of Palestinian strains (13/19) along with those from China; confirming low level of genetic diversity present in this cluster compared to clusters II and III. Tajima’s D and Fu-Li’sF tests were negative in cluster I and showing statistically significant departure from neutrality (random mutation) (P<0.01). As for CVB5 genotype, cluster III recorded the lowest genetic diversity indices and neutrality tests with negative but insignificant values (Table 3 and 4). Both E18 and CVB5 showed a combination of high haplotype diversity (Hd), but low genetic diversity (π). Also, both showed overall negative values of neutrality tests; Tajima’s D and Fu-Li’s F.

**Table 3.** Haplotype-nucleotide diversity and neutrality tests of three cluster of E18 as calculated for the VP1 gene.

| Cluster | Haplotype- nucleotide diversity |  |  |  |  |  |  | Neutrality tests |  |
| --- | --- | --- | --- | --- | --- | --- | --- | --- | --- |
| | n | h | h:n | Hd±SD | $\pi \pm$SD | K | S | Tajima's D | Fu-Li's F |
| RED-I | 19 | 12 | 0.6:1 | 0.92± 0.05 | 0.03± 0.01 | 11.3 | 67 | -1.80* | -2.97* |
| Green-II | 7 | 7 | 1:1 | 1.0± 0.08 | 0.07± 0.01 | 61.9 | 165 | -0.57 | -0.56 |
| Blue-III | 10 | 9 | 0.9:1 | 1.0± 0.05 | 0.06± 0.01 | 55.2 | 171 | -0.93 | -1.12 |
| **Total** | **41** | **34** | **0.9:1** | **0.98± 0.01** | **0.12± 0.02** | **35.5** | **184** | **-1.50** | **-2.80** |

n: Number of sequences, h: Number of Haplotypes, Hd: Haplotype (gene) diversity, $\pi$ : Nucleotide diversity (per site), K: Average number of nucleotide differences between two randomly chosen sequences from within in the population, S: Number of variable/segregating sites. (1 outgroup, 4 have no clear cluster). \*: $P < 0.01$ .

**Table 4.** Haplotype/nucleotide diversity and neutrality tests of three cluster of CVB5 as calculated for the VP1 gene.

| Cluster | Haplotype nucleotide diversity |  |  |  |  |  |  | Neutrality tests |  |
| --- | --- | --- | --- | --- | --- | --- | --- | --- | --- |
| | n | h | n:h | Hd±SD | $\pi \pm$SD | K | S | Tajima's D | Fu-Li's F |
| Blue-I | 14 | 14 | 1:1 | 1.0± 0.03 | 0.13± 0.009 | 89.3 | 89 | 0.05 | 0.36 |
| Green-II | 13 | 13 | 1:1 | 1.0± 0.03 | 0.10± 0.01 | 28.3 | 74 | -0.257 | -0.27 |
| RED-III | 19 | 18 | 1:1 | 0.94± 0.02 | 0.08± 0.007 | 24.3 | 75 | -0.55 | -1.13 |
| **Total** | **46** | **45** | **1:1** | **0.99± 0.006** | **0.17± 0.01** | **41.04** | **133** | **-0.75** | **-1.76** |

n: Number of sequences, h: Number of Haplotypes, Hd: Haplotype (gene) diversity, $\pi$ : Nucleotide diversity (per site), K: Average number of nucleotide differences between two randomly chosen sequences from within in the population, S: Number of variable/segregating sites. (1 outgroup, 4 have no clear cluster)

Inter-population pairwise genetic distance (Fst) in the three E18 populations ranged from 0.39 to 0.63 with Nm value from 0.29 to 0.78 (Table 5 and 6). Fst for cluster I containing all Palestinian strains compared to clusters II and III were high (0.51 and 0.63) (positive Tajima’s D) indicating population differentiation with low gene flow, Nm (0.48 and 0.29). However, genetic differentiation between clusters II and III is low (Fst=0.39) with high Nm (0.78), but still reflected genetic differentiation. This is supported by the negative value of Gst (-0.006).

**Table 5:**
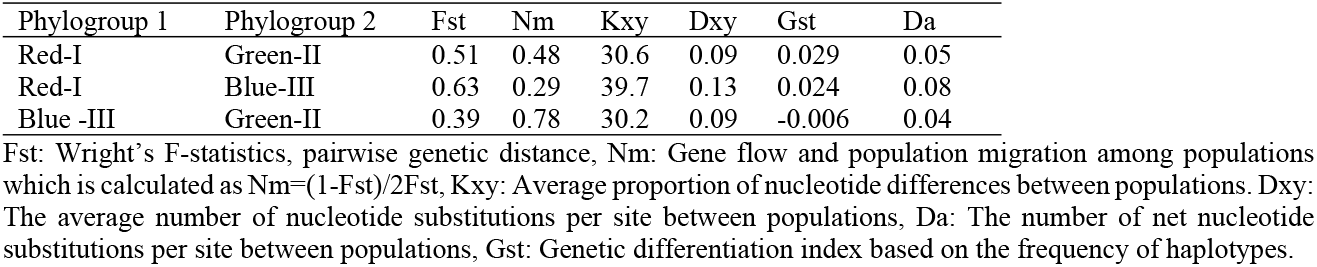
Gene flow and genetic differentiation indices between the three E18 clusters estimated from VP1 sequences.

**Table 6:** Gene flow and genetic differentiation indices between the three CB5 clusters estimated from VP1 sequences.

| Phylogroup 1 | Phylogroup 2 | $F_{st}$ | $N_m$ | $K_{xy}$ | $D_{xy}$ | $G_{st}$ | $D_a$ |
| --- | --- | --- | --- | --- | --- | --- | --- |
| Red-III | Green-II | 0.54 | 0.43 | 48.8 | 0.19 | 0.005 | 0.11 |
| Red-III | Blue-I | 0.49 | 0.52 | 51.8 | 0.21 | 0.005 | 0.10 |
| Blue-I | Green-II | 0.32 | 1.06 | 43.9 | 0.18 | 0.00005 | 0.06 |

$F_{st}$ : Wright's F-statistics, pairwise genetic distance, $N_m$ : Gene flow and population migration among populations which is calculated as $N_m=(1-F_{st})/2F_{st}$ , $K_{xy}$ : Average proportion of nucleotide differences between populations $D_{xy}$ : The average number of nucleotide substitutions per site between populations, $D_a$ : The number of net nucleotide substitutions per site between populations, $G_{st}$ : Genetic differentiation index based on the frequency of haplotypes.

As for CVB5, cluster III which contained most of the Palestinian strains is differentiated from the other two clusters, I and II, as reflected by the high Fst and low Nm values (Fst=0.54, Nm=0.43) and (Fst=0.49 and 0.52), respectively. At the same time, the genetic differentiation between clusters I and II is relatively low as supported by very low Gst value (Gst=0.00005) and relatively low Fst (0.32) with high gene flow (Nm=1.06).

## Discussion

So far, only one report in Palestine confirmed the isolation of 7 different enteroviruses genotypes including echovirus 13, 14, 9, 30, 16, 6 and coxsackievirus B5 from sporadic cases of aseptic meningitis and/or sepsis like illness[10]. In the present study, all of the HEV positive cases occurred in children less than 7 years old and 50% of them < 1-year-old. Highest rate of enterovirus-positive cases was reported previously in children less than 1 year of age in Palestine, United States, and Korea [10, 38]. On the contrary, other studies, reported that HEV aseptic meningitis occurs most frequently in patients with age range of 3–12 years’ old [13-15, 39]. The discrepancy in the HEV age groups could be due to the source of cases whether sporadic or from an outbreak.

The present study showed that E18 and CVB5 were the most predominant genotypes, representing, 50% and 34.6% respectively. Several recent studies in Germany, Taiwan, Korea, and China reported that both E18 and CVB5 were the most predominant HEVs reported from sporadic and outbreak cases of aseptic meningitis [13, 15-16, 40].

All E18 sequences included in the phylogenetic analysis grouped into three major genetic clusters. The E18 isolates from Palestine clustered together along with those from China reported in 2015 in cluster I. Clusters II and III contain E18 from Australia, France, USA, Sweden, Russian, Germany and Netherlands reported in the periods 2005-2012 and 1999-2012, respectively. Other minor clusters included cluster IV and V which included the prototype Metacalf and few HEV reported from India and China in the period 2000-2008. HEV types in clusters IV and V may have circulated during a limited period and then disappeared. Similar results reported a genetic divergence in the complete VP1 gene of the E18 that resulted in the formation of five phylogenetic clusters [40]. These viruses were reported recently from China, India, South Korea, Australia,

Netherlands, Germany, Sweden, Russia and France [40]. Phylogenetic analysis of partial or complete VP1 gene of E11, E30, E13, and E6 of the HEV-B species revealed high genetic diversity showing several clusters and sub-clusters [40]. Accordingly, such data indicate that the same HEV genotype may circulate in different geographies at different times. Therefore, and due to lack of reporting HEVs in Palestine, the possibility of knowing whether E18 isolates in cluster I are recently introduced or have already been circulating before remains a dilemma.

The CVB5 sequences grouped into three genetic clusters (Fig 1). Cluster III contained the 9 Palestinian strains isolated in 2017 along with CVB5 from Brazil, France, Cyprus, Germany and Australia reported in the period from 2006 to 2017. The 2 Palestinian isolates in 2013 grouped with a more recent CVB5 in cluster II (2003-2014) from China, Denmark, France and Netherland. Cluster I contained the prototype Faulkner and isolates from France, Germany, Japan, Korea, China, Hong Kong, India, Finland and Sweden. Recently, few studies compared the partial or the complete VP1 gene revealing genetic diversity of CVB5 in 2-5 clusters [41-43]. Accordingly, the findings in the present study reaffirm that two or more clusters may co-circulate and coevolve in the same region as a result of the genetic diversity forces such as mutation or recombination.

The recombination for both species E18 and CVB5 was minimal, if at all present between the study sequences which could be explained by the infrequency of recombination within the capsid region, VP1 [19-20]. This puts forward mutation rates as the main cause of genetic diversity. The significant departure from neutrality as confirmed by negative Tajima’s D and Fu-Li’s F accompanied by high haplotype (Hd) and low nucleotide diversity (π) for cluster I of E18 (Table3) may suggest recent rapid population expansion phase following bottleneck or genetic hitchhiking (genetic draft or gene sweep) that bring excess number of rare alleles. This was supported by the negative values Tajima’s D and Fu-Li’s F tests in the individual populations and the overall value for the three populations. A study in Taiwan showed that partial VP1 genes of E18 have low genetic diversity with high similarity between regions [14]. The clustering of the Palestinian and Chinese isolates in cluster I could have been brought about by the activity of Palestinian traders to China and back. Cluster II and III had similar diversity indices, but without any significance.

Cluster III of CVB5 showed lower genetic diversity than the others (Table 4), yet insignificant; suggesting that cluster III may have undergone a neutral or contraction period or may be due to small subpopulations with limited number of sequences to yield sufficient statistical power. A study from Korea revealed very high similarity in the partial VP1 sequences among Korean samples (intra-population similarity) and between Chinese strains (inter-population similarity)[16]. The genetic similarity of the Palestinian, Korean and Taiwanese E18 and CVB5 strains to those from China, may hint to their dispersal from China to the rest of the world [14, 16]. However, a more extensive study of several endemic areas should be conducted to confirm the ancestral origin of E18 and CVB5, the route of spread and the distribution.

The estimations of inter-population comparison indices (Fst, Nm, Kxy, Dxy, Gst and Da) (Tables5 and 6) support high level of genetic differentiation between the three main clusters of both E18 and CVB5. These results are supportive of the cluster in phylogentic analyses (Fig1).

In conclusion, the present study unravels three main clusters for each of the E18 and CVB5 with both showing high haplotype diversity compared to lower genetic diversity. The Palestinian isolates grouped mainly into one cluster for each viral species. Finally, present study supports close genetic relationship between Palestinian HEV species as confirmed by population genetics and phylogenetic analyses.

